# *Cryptococcus neoformans* evades pulmonary immunity by modulating xylose precursor transport

**DOI:** 10.1101/741017

**Authors:** Lucy X. Li, Camaron R. Hole, Javier Rangel-Moreno, Shabaana A. Khader, Tamara L. Doering

**Affiliations:** Department of Molecular Microbiology, Washington University School of Medicine, St. Louis, Missouri, USA; Department of Medicine, Allergy/Immunology, and Rheumatology, University of Rochester School of Medicine, Rochester, New York, USA

## Abstract

*Cryptococcus neoformans* is a fungal pathogen that kills almost 200,000 people each year and is distinguished by abundant and unique surface glycan structures that are rich in xylose. A mutant strain of *C. neoformans* that cannot transport xylose precursors into the secretory compartment is severely attenuated in virulence in mice, yet surprisingly is not cleared. We found that this strain failed to induce the non-protective T helper cell type 2 (Th2) responses characteristic of wild-type infection, instead promoting sustained Interleukin (IL)-12p40 induction and increased IL-17A (IL-17) production. It also stimulated dendritic cells to release high levels of pro-inflammatory cytokines, a behavior we linked to xylose expression. We further discovered that inducible bronchus associated lymphoid tissue (iBALT) forms in response to infection with either wild-type cryptococci or the mutant strain with reduced surface xylose; although iBALT formation is slowed in the latter case, the tissue is better organized. Finally, our temporal studies suggest that lymphoid structures in the lung restrict the spread of mutant fungi for at least 18 weeks after infection, in contrast to ineffective control of the pathogen after infection with wild-type cells. These studies demonstrate the role of xylose in modulation of host response to a fungal pathogen and show that cryptococcal infection triggers iBALT formation.

## Introduction

*Cryptococcus neoformans* is a ubiquitous environmental fungus that causes pneumonia and meningitis. This opportunistic pathogen infects over a million individuals each year, with overall mortality exceeding 20% (1-3). Patient immune status is the main determinant of infection outcome, highlighting the importance of efficient host immune responses in the control of cryptococcosis.

*C. neoformans* infection begins when the organism is inhaled, which is followed by its proliferation in the lungs. This pulmonary infection may then disseminate to the brain, where it causes a frequently lethal meningoencephalitis. In the lungs, *C. neoformans* first interacts with host macrophages and dendritic cells (DC). These engulf the fungus and migrate to draining lymph nodes, where they present antigen to T cells to initiate the adaptive immune response. The protective immune response to this infection is primarily mediated by T cells: T helper cell type 1 (Th1) responses are protective against *C. neoformans* (4) and T helper type 17 (Th17) responses likely participate in pathogen clearance at mucosal surfaces (5-7). In contrast, T helper cell type 2 (Th2) responses are considered detrimental for protective immunity, leading to fungal growth and facilitating dissemination to the CNS (8, 9). The induction of these non-protective responses is not well understood, but cryptococcal glycans may contribute to this process (10).

Glycan structures participate in antigen recognition, immune activation, and immune regulation in multiple organisms (11). In *C. neoformans* the major virulence factor, a polysaccharide capsule, modulates the immune response via multiple mechanisms, including dampening immune cell activation and impeding phagocytosis (12, 13). The capsule is primarily composed of two large polysaccharides: glucuronoxylomannan (GXM) and glucuronoxylomannogalactan (GXMGal) (14, 15). One major component of both polymers is xylose (Xyl), which comprises almost one fourth of the polysaccharide capsule mass (16). Xylose also occurs in cryptococcal glycolipids (17) and as both Xyl and Xyl-phosphate modifications of protein-linked glycans (18-20). We demonstrate here that Xyl plays a critical role in the immune recognition of, and response to, *C. neoformans.*

The incorporation of Xyl into cryptococcal glycan structures occurs in the lumen of the secretory pathway, via enzymatic reactions that use the substrate molecule UDP-Xyl. This xylose donor is imported into the synthetic compartment by two transporters, Uxt1 and Uxt2 (21). A mutant strain that lacks both transporters (*uxt1*Δ *uxt2*Δ) exhibits defects in xylosylation, including incomplete synthesis of capsule polysaccharides, and considerable attenuation of virulence in mouse models of infection. Surprisingly, despite its compromised virulence, which contrasts with the rapid lethality of wild-type (WT) infection, the mutant strain persists for months in the lungs of infected mice (21).

In pursuing the mechanism of *uxt1*Δ *uxt2*Δ persistence, we found that infection with the mutant failed to induce the Th2 responses provoked by WT fungi, although it did promote production of IL-12p40 and IL-17, likely via enhanced stimulation of DCs. The corresponding inflammatory response was enriched in T and B cells, which were required for the prolonged survival of the infected animals. Inflammation in the lung, although it increased slowly compared to the response to WT infection, appeared to effectively contain the mutant yeast cells for an extended period after infection. Finally, we observed the formation of inducible bronchus associated lymphoid tissue (iBALT) in the lungs of both WT and *uxt1*Δ *uxt2*Δ infected mice. iBALT resembles secondary lymphoid organs (lymph nodes) in cellular composition and organization, with two distinctive zones: a central B cell follicle and surrounding T cell zone (22). The T cell zone contains CD4 and CD8 T cells as well as DCs (23); the B cell zone contains mainly follicular B cells and its organization is maintained by the continuous production of CXCL13 (24). DC activation and cytokine production are also required for iBALT formation and maintenance (23, 25). In the lungs of mice infected with the *uxt1*Δ *uxt2*Δ mutant, iBALT formation was significantly delayed. However, once formed, it was better organized than the iBALT produced when mice were infected with fully xylosylated WT *C. neoformans.* Together, these results suggest that luminal Xyl modifications of cryptococcal glycoconjugates alter immune recognition and activation of immune cells, events that are required for efficient control of *C. neoformans* infection.

## Results

### UDP-Xyl transport is required for cryptococcal virulence and dissemination

We previously observed that a *C. neoformans* strain lacking UDP-Xyl transporters (*uxt1*Δ *uxt2*Δ) was highly attenuated in virulence compared to the WT parental strain KN99α, yet remained in the lungs of asymptomatic mice for at least 100 days after infection (21). While defects in the mutant strain, such as altered capsule and a slightly decreased growth rate, might explain its reduced virulence, this persistence was surprising. To investigate the development of this chronic infection, we first examined the kinetics of fungal burden in mice after intranasal inoculation with WT and mutant strains. In A/JCr mice infected intranasally with WT fungi, which typically succumb to infection roughly 18 days post-infection (dpi), we observed a rapid and significant increase in pulmonary fungal burden (Fig. 1A) along with significant dissemination to the spleen (Fig. 1B) and brain (Fig. 1C). In contrast, the pulmonary burden in *uxt1*Δ *uxt2*Δ-infected mice increased slowly from initial inoculum levels to a modest peak at 63 dpi (Fig. 1A). Colony-forming units in the lung then gradually declined, although even after six months only 30% of mice showed complete clearance (Fig. 1A, 189 dpi). Although *uxt1*Δ *uxt2*Δ infection was not restricted to the lungs, only a fraction of the mice had measurable mutant cells in the spleen (Fig. 1B) and brain (Fig. 1C) at 63 dpi. Furthermore, this limited dissemination was only observed at the times of highest lung burden (63 and 126 dpi), with no fungi detected at distal sites by 189 dpi (Fig. 1B, Fig. 1C).

**Figure 1.**
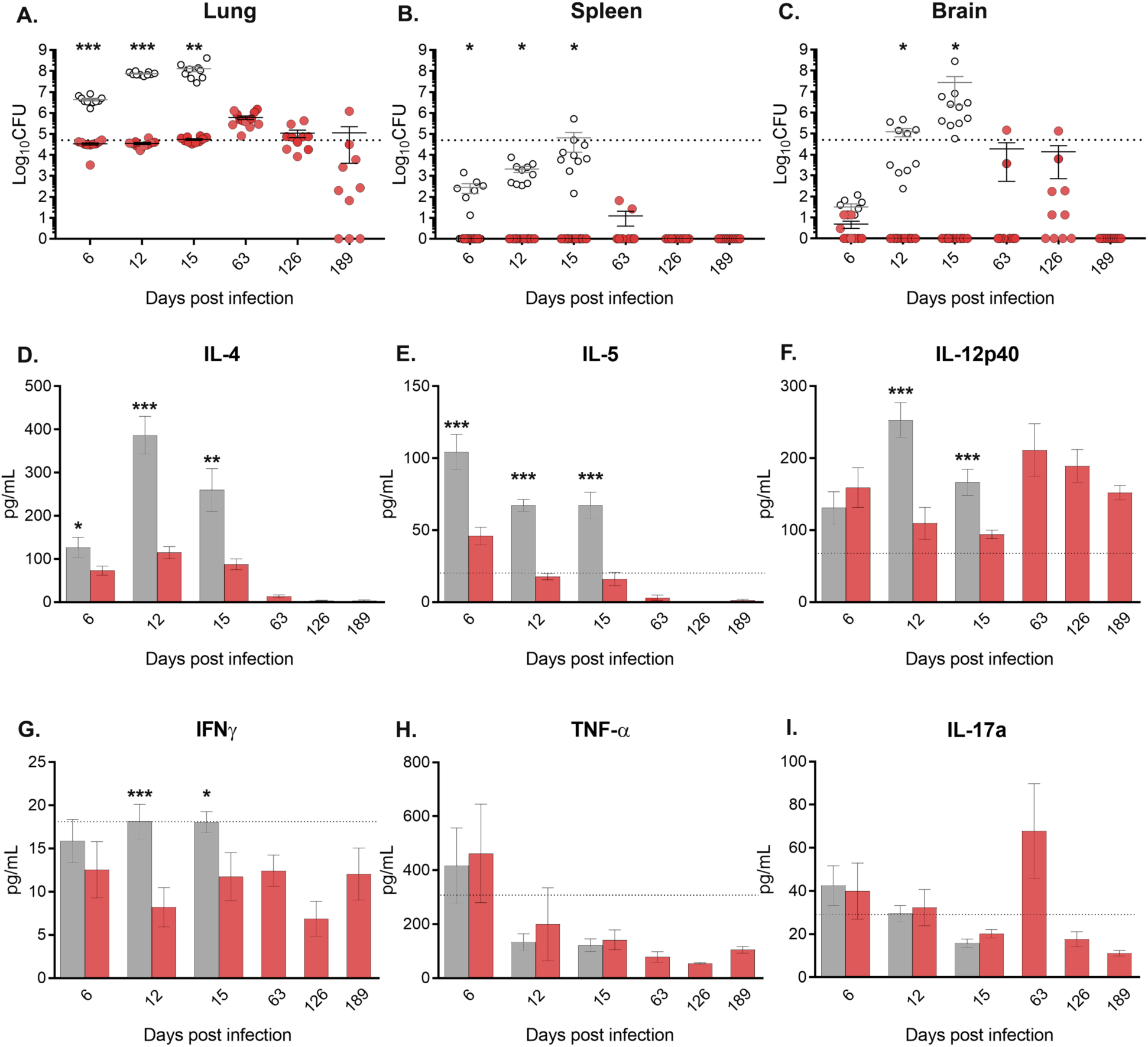
Fungal burden and pulmonary cytokine production in WT and *uxt1*Δ *uxt2*Δ mice. All panels show the combined results of two independent experiments (5 mice per group per experiment). (A-C) Tissue homogenates of infected A/JCr mice were plated for CFUs at the indicated dpi (open symbols, WT; red symbols, *uxt1*Δ *uxt2*Δ; dashed line, initial inoculum). Values from individual mice are plotted along with the mean ± SEM. (D-I) Cytokine levels in lung homogenates at the indicated dpi (dashed line, naïve; gray bars, WT; red bars, *uxt1*Δ *uxt2*Δ). *, *p* < 0.05; **, *p* < 0.01; ***, *p <* 0.005 for Student’s t*-*test comparing WT to mutant.

### Cryptococcal xylosylation influences production of pulmonary cytokines and cellular immune response

We next sought to identify the mechanism(s) responsible for the unexpected persistence of *uxt1*Δ *uxt2*Δ after intranasal infection. As summarized above, protection against *C. neoformans* is generally associated with production of type 1 cytokines like interleukin (IL)-2, IL-12, interferon gamma (IFN-γ), and tumor necrosis factor alpha (TNF-α). In contrast, a Th2 response to cryptococcal infection (characterized by the production of IL-4, IL-5, and IL-13) compromises control of pulmonary fungal burden (26) and facilitates dissemination to the central nervous system (4, 26).

We hypothesized that the protracted host clearance of the *uxt1*Δ *uxt2*Δ mutant reflected an altered host immune response to cells that display reduced xylose. To assess the host response, we first analyzed cytokine levels over time in lung homogenates from mice infected with WT or *uxt1*Δ *uxt2*Δ strains (Fig. 1). In mice infected with WT cryptococci, we observed a strong Th2-type response, with significant induction of IL-4 and IL-5, over the 15 days prior to sacrifice (Fig. 1D, Fig. 1E). Other proinflammatory cytokines (IL-1α, IL1β, IL-6) were modestly increased (Supp. Fig. 1). In contrast, *uxt1*Δ *uxt2*Δ cells elicited a more quiescent Th2 cytokine response at early stages of infection. Furthermore, at the peak of fungal burden in *uxt1*Δ *uxt2*Δ-infected mice (day 63, Fig. 1A), there was a significant increase in the Th1 polarizing cytokine IL-12p40 compared to the levels observed in naïve lungs (Fig. 1F); IL-12p40 levels then slowly declined, along with fungal burden. Although the levels of IFNγ and TNFα in mutant-infected mice resembled those in the lungs of naïve mice (Fig. 1G, Fig. 1H), IL-17 trended higher at the peak of infection (Fig. 1I). Overall, while WT infection rapidly induced non-protective Th2 responses, infection with *uxt1*Δ *uxt2*Δ failed to do so (even at the peak of fungal burden) and instead promoted the sustained induction of IL-12p40 and increased production of IL-17.

To measure the host adaptive immune response to *uxt1*Δ *uxt2*Δ, we used flow cytometry to count lymphocytes in the lungs of mice infected with WT or *uxt1*Δ *uxt2*Δ cells. By 15 dpi, the B cell fraction of viable leukocytes in mutant-infected mice significantly exceeded the value for WT-infected mice (Fig. 2A); this fraction further increased as mutant infection progressed, to peak at over 30%. The same pattern was seen for CD4^+^ and CD8^+^ T cells (Fig. 2B, Fig. 2C), initiating even earlier (at 12 dpi) and remaining high throughout the time course. The total numbers of cells in all three populations remained elevated through 126 dpi (Fig. 2A-C, right), although they subsequently declined.

**Figure 2.**
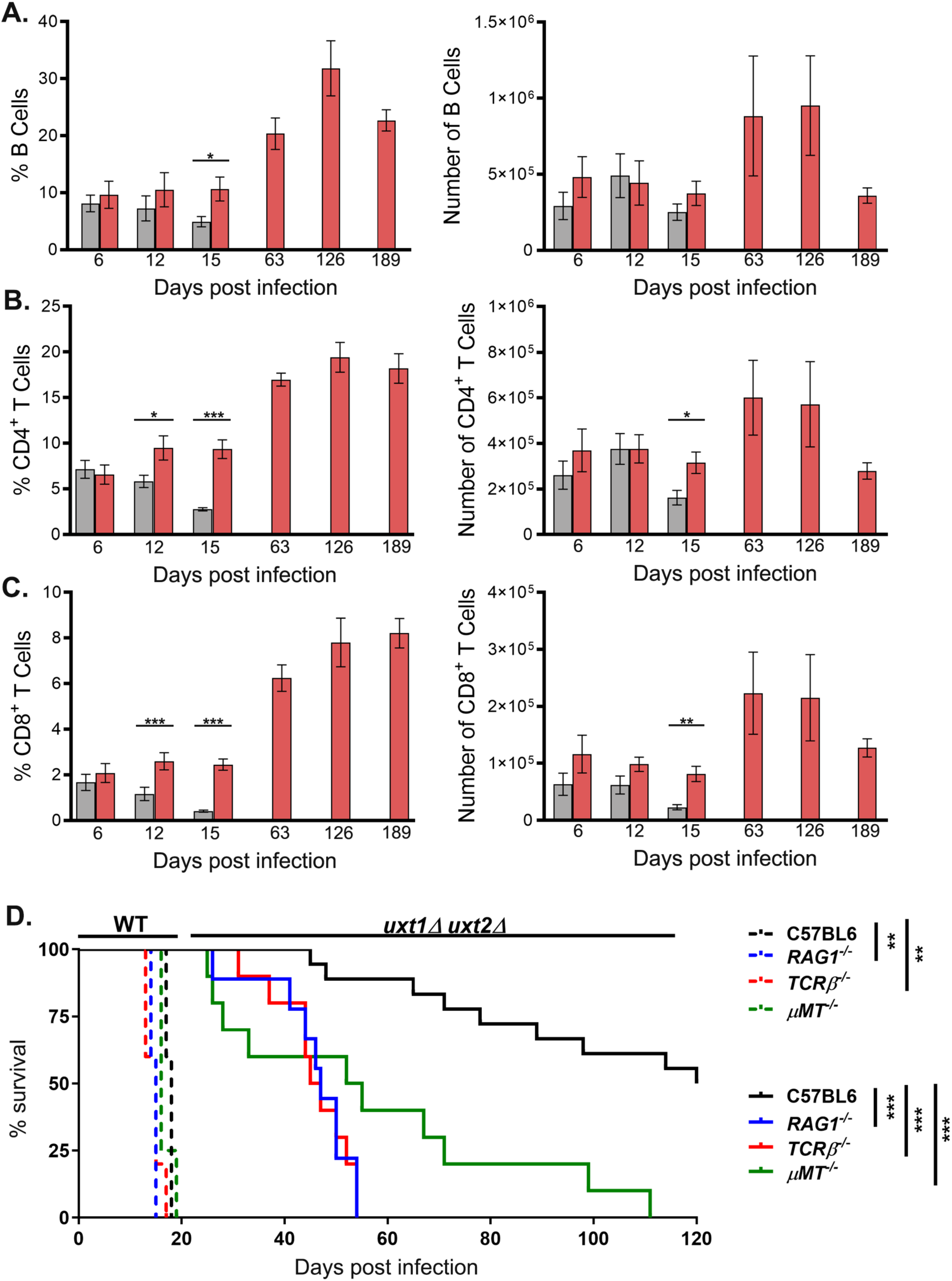
Accumulation of T and B cells in the lungs of mice infected with *uxt1*Δ *uxt2*Δ cells is required to prevent disease progression. CD19^+^ B cells (A), CD4^+^ T cells (B), and CD8^+^ T cells (C) were enumerated in the lungs of infected mice by flow cytometry (gray, WT; red, *uxt1*Δ *uxt2*Δ). Plotted are the combined mean ± SEM values from two independent experiments (5 mice per group per experiment). *, *p* < 0.05; **, *p* < 0.01; ***, *p* < 0.005 by Student’s t-test. (D) Survival of mice after intranasal inoculation with 5 × 10^4^ cells of WT (n = 5) or *uxt1*Δ *uxt2*Δ (n = 10). Results shown are combined from two independent experiments. **, *p* < 0.01; ***, *p* < 0.005 by the log-rank test.

### Survival of uxt1Δ uxt2Δ-infected mice is dependent on T cells

Because our results suggested that T and B cells play a protective role that contributes to the prolonged survival of *uxt1*Δ *uxt2*Δ-infected mice, we next examined whether ablation of these cell types altered the course of infection. Because the mouse strains used for these studies were engineered in the C57BL/6 background, we included infections of this strain as controls. C57BL/6 mice are less resistant to *C. neoformans* than the A/JCr mice used above (27), as shown by their eventual susceptibility to *uxt1*Δ *uxt2*Δ infection (Fig. 2D, solid black line). Notably, *Rag1*-/- mice in this background were significantly more susceptible to the mutant fungi: all succumbed by 55 dpi (Fig 2D; solid blue line). These animals still survived longer than WT-infected mice, possibly due to the slightly slower growth of the mutant which we previously observed (21).

Since *Rag1*-/- mice lack both T and B cells, we also infected mice deficient in either T cells (*TCRβ* -/-) or B cells (*μMT*) with *uxt1*Δ *uxt2*Δ cryptococci. B cell deficient mice exhibited a more protracted course of disease than *Rag1*-/- animals, succumbing over an 85-day period (Fig. 2D, solid green line), while mice lacking only T cells succumbed to infection with kinetics similar to those of the *Rag1*-/- mice (Fig. 2D, solid red line). In contrast, all mice inoculated with WT *C. neoformans* succumbed by 20 dpi (Fig. 2D; dashed lines). Of this group, *Rag1*-/- and *TCRβ*-/- mice still succumbed slightly faster than C57BL/6 and μMT mice (Fig. 2D). Together, these data show that T cells are the prominent cell type responsible for increased survival of *uxt1*Δ *uxt2*Δ-infected mice, although B cells may play a lesser role in protection.

### Reduced surface xylose of C. neoformans stimulates DC activation

The antigen-driven activation of DCs is critical to induce protective immunity against *C. neoformans* (28) and it is likely that their consequent cytokine and chemokine production are responsible for T cell recruitment (23). We speculated that an enhanced DC response to xylose-deficient fungi could explain the rise in T cells during *uxt1*Δ *uxt2*Δ infection (Fig. 2), even in the absence of expanded numbers of DCs (Fig. 3A). To test this idea, we grew bone marrow derived DCs, stimulated them for 24 hours with heat-killed *C. neoformans*, and measured the production of pro-inflammatory cytokines. In these studies we used WT fungi, which display antigens modified with Xyl, or *uxt1*Δ *uxt2*Δ fungi, which are impaired in these modifications. We also tested strains with individual *uxt* mutations: *uxt1*Δ shows an intermediate level of Xyl utilization while *uxt2*Δ resembles WT (21). Finally, we assayed *uxs1*Δ cells, which lack all Xyl modification because they cannot synthesize UDP-Xyl (29), and an acapsular control strain (*cap59*Δ) known to induce a potent DC response (30).

**Figure 3.**
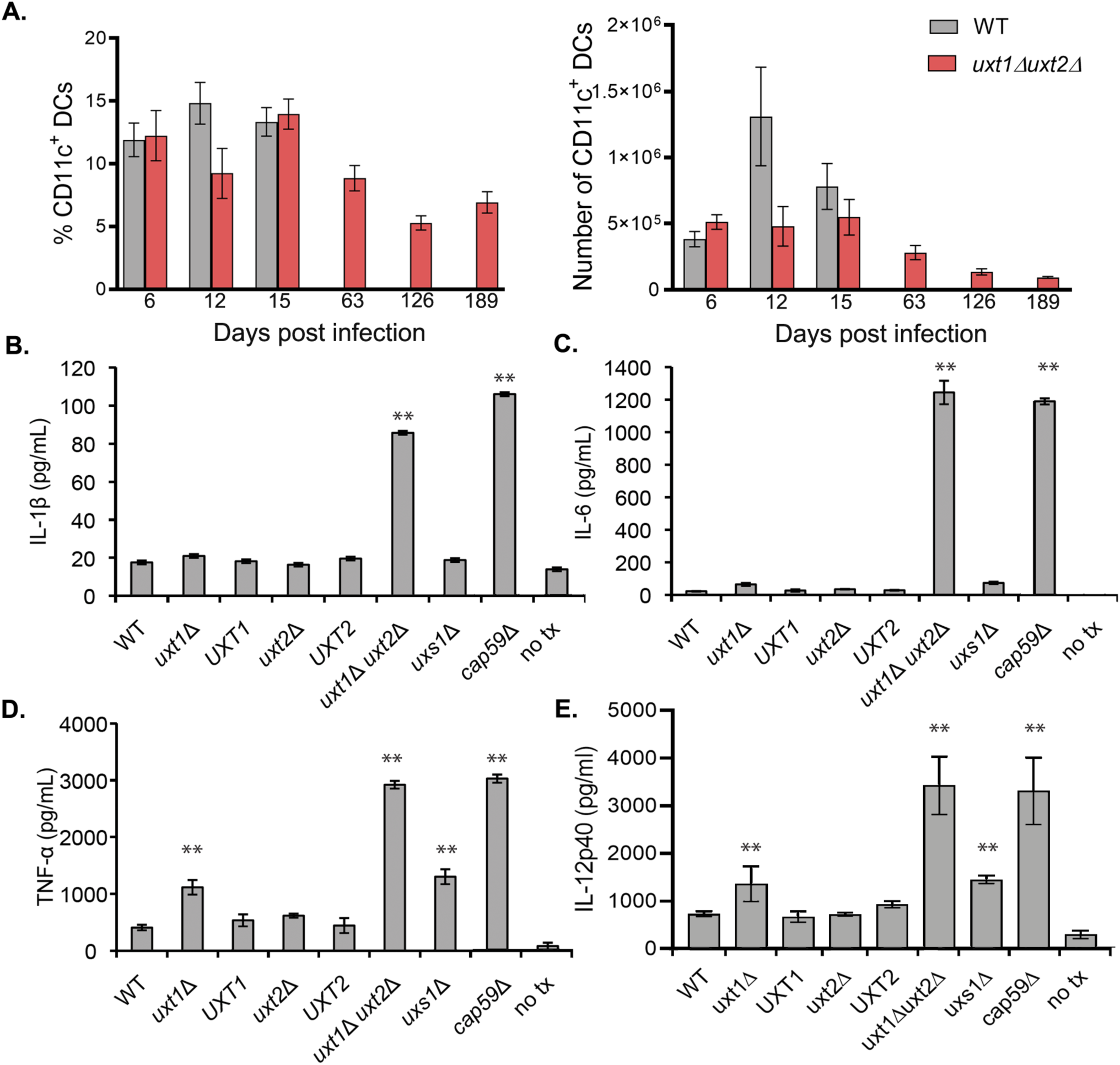
Cytokine production by dendritic cells stimulated with cryptococcal antigens. (A) CD11c+ DC populations in the lungs of infected mice were quantified by flow analysis. (B-E) DCs were co-incubated for 24 h with heat-killed cells of the strains indicated or subjected to no treatment (no tx) and levels of IL-1β (B), IL-6 (C), TNF-α (D), and IL-12p40 (E) in the supernatant were quantified by ELISA. Shown are the mean ± SD (n = 3) values for one representative experiment from five independent experiments that yielded similar results. **, *p* < 0.01 by one-way ANOVA with Tukey’s post hoc test.

We found that bone marrow derived DCs (BMDCs) cultured with *uxt1*Δ *uxt2*Δ cells released high levels of IL-1β and IL-6, comparable to the levels observed following exposure to a completely acapsular control strain, *cap59*Δ (Fig. 3B-C). In contrast, individual *uxt* mutants and *uxs1*Δ induced IL-1β and IL-6 levels similar to those induced by WT fungi (close to background levels; Fig. 3B-C). TNF-α and IL-12p40 similarly showed the greatest response upon challenge with *uxt1*Δ *uxt2*Δ, with levels like those induced by the acapsular strain, although a slight increase was also noted with other xylose-deficient strains (*uxt1*Δ and *uxs1*Δ; Fig. 3D-E)). These results indicate that early interaction with DCs and consequent induction of proinflammatory cytokines are influenced by the surface Xyl ex180 pression of *C. neoformans*.

### The cellular response to cryptococcal infection includes the development of inducible bronchus-associated lymphoid tissue (iBALT)

Our data suggested that *uxt1*Δ *uxt2*Δ mutant cells stimulate elevated production of proinflammatory cytokines by DCs, which in turn significantly increases the T and B cells in infected lungs (in both absolute numbers and as a fraction of total leukocytes). Flow cytometry, however, provides no information as to the distribution of these cell populations within the lung. To futher probe the cellular immune response, we therefore examined lung histology, focusing on late time points of infection: 15 dpi for mice infected with WT fungi and 126 and 189 dpi for *uxt1*Δ *uxt2*Δ-infected mice (also including 15 dpi to allow direct comparison to WT). We used Movat’s stain for these studies (Fig. 4), because the Alcian Blue component stains cryptococcal cell walls and capsules turquoise, allowing visualization of fungi as well as of host cells (31). In WT-infected mice at 15 dpi we saw the typical pattern of overwhelming fungal burden throughout the lung, with little organized inflammation (32). The tissue architecture was largely destroyed and there was a preponderance of enlarged fungi with thick capsules. In contrast, mutant-infected mice at the same time point showed relatively preserved tissue architecture, with limited regions of organized inflammation. Most cryptococci were restricted to these areas and overall fungal burden was noticeably lower, as expected for this mutant (Fig. 1A). By 126 dpi of infection with mutant fungi, the regions of inflammation were larger and more highly consolidated; fungi were rarely observed in other areas of the lung. The last time point of mutant infection (189 dpi) looked similar, although inflammation appeared slightly reduced, consistent with the decrease in B and T cells that we observed at that time point (Fig. 2). Lymphoid structures were also less dense; fungi were less restricted to the cen205 ters of these regions and occasionally appeared in lung tissue beyond them.

**Figure 4.**
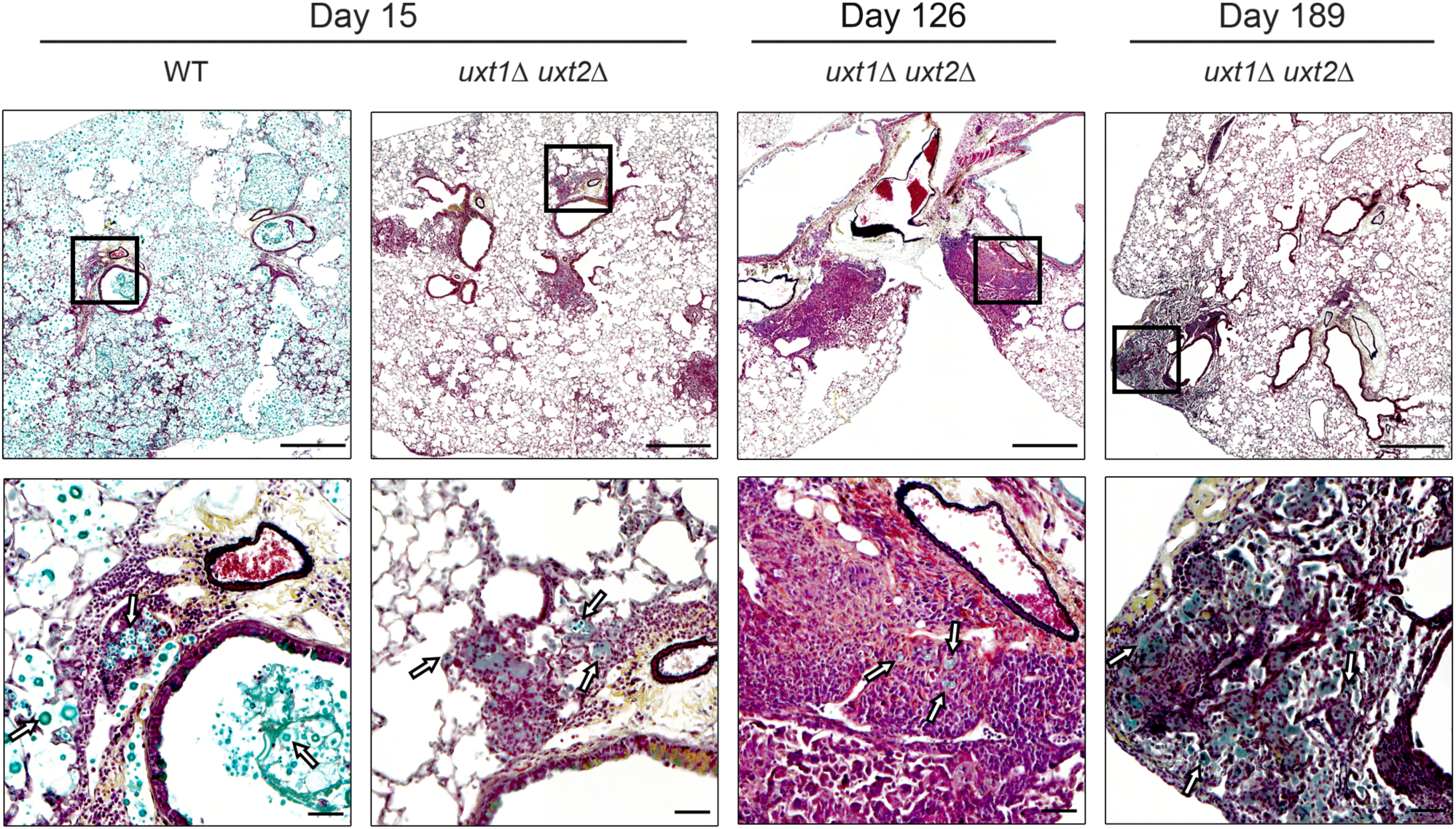
Mutant cryptococci are primarily restricted to lymphoid structures. Micrographs show Movat-stained lung sections from A/JCr mice infected with WT or *uxt1Δuxt2Δ* as above. Scale bars, 100 μm (top row) and 10 μm (bottom row). White arrows highlight fungi (stained turquoise). Images shown are representative of two independent studies (4-5 mice per group per experiment).

In both groups of mice (WT- and mutant-infected), we noted distinctive lymphocytic accumulations proximal to the basal side of the bronchial epithelium (Fig. 4). To probe these structures, we stained lung sections for T cells (CD3), B cells (B220), and plasma cells (IgG). In all samples we observed B cell follicles surrounded by CD3^+^ T cell cuffs (Fig 5A), suggestive of inducible bronchus-associated lymphoid tissue (iBALT) (24). Consistent with identification as iBALT, these structures stained positive for CXCL13 (Fig. 5B).

**Figure 5.**
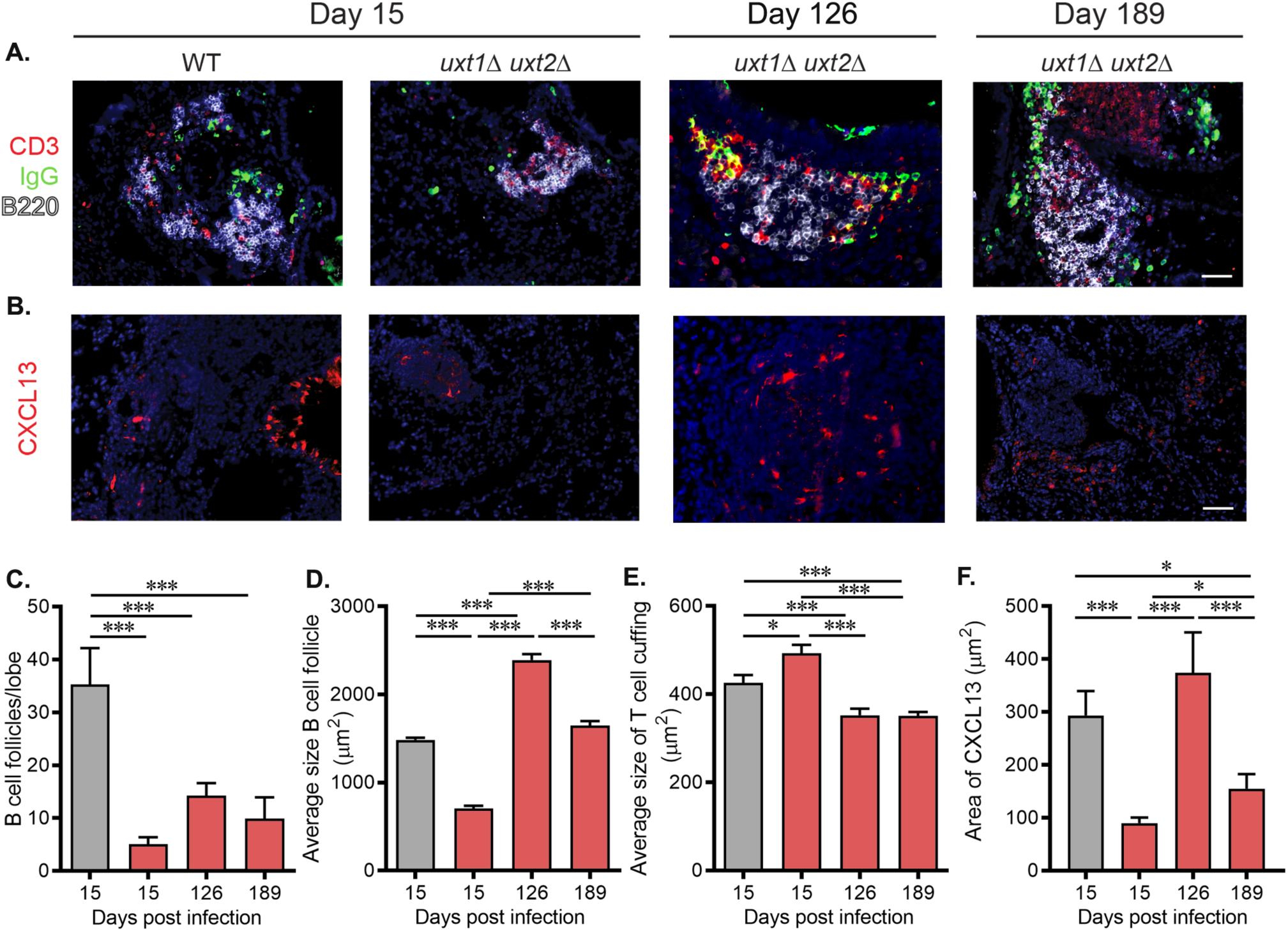
iBALT complexity in the context of *C. neoformans* infection. (A-B) Representative immunofluorescent staining of B cells (B220^+^), T cells (CD3^+^), plasma cells (IgG^+^), and CXCL13 in the lungs of infected A/JCr mice. Scale bar = 100 μm. (C-E) Morphometric analysis of iBALT structures in A/JCr mice infected with WT (gray bars) and *uxt1*Δ *uxt2*Δ (red bars). (F) Quantification of immunofluorescent staining of CXCL13. Plots show the combined mean ± SEM of two independent studies (4-5 mice per group per experiment); colors as in panels C-E. *, *p* < 0.05; ***, *p* < 0.005 by Student’s t-test.

To examine iBALT development, we performed blinded morphometric analysis of lung tissue from mice infected with WT and mutant cryptococci. At 15 dpi with WT fungi, we observed numerous iBALT structures, with compact B cell follicles surrounded by T cells (Figs. 5C-E). In contrast, B cell follicles were considerably fewer and less organized at the same time point in mice infected with the *uxt1*Δ *uxt2*Δ strain (Figs. 5C-E). The limited formation of iBALT structures was also characterized by increased peri-vascular accumulation of T cells (Fig. 5A) and decreased production of CXCL13 (Fig. 5B, Fig. 5F). By 126 dpi with *uxt1*Δ *uxt2*Δ, however, although iBALT structures were still less abundant than those found in WT infection (Fig. 5C), their average B cell follicle size was significantly larger (Fig. 5D). Additionally, the *uxt1*Δ *uxt2*Δ iBALT was more organized, with reduced T cell cuffing (Fig. 5E), increased CXCL13 (Fig. 5F) and more IgG^+^ plasma cells (Fig. 5A). By 189 dpi, these structures regressed slightly, consistent with the reduced lymphocyte infiltration, cytokine levels, and fungal burden at this time point. WT cryptococci thus induce robust early formation of iBALT, while the *uxt1*Δ *uxt2*Δ mutant induces delayed but ultimately larger and better organized iBALT.

## Discussion

We wished to investigate the unexpected observation that a mutant strain of *C. neoformans*, despite significant defects in surface glycoconjugates, nevertheless persisted for many months in infected mice. We found that *uxt1*Δ *uxt2*Δ cells showed an initial delay after intranasal infection, but then began to accumulate slowly, eventually rising to a modest peak, at which time (63 dpi) we also observed limited dissemination. The lung burden then began to gradually decline, although by the end point of the study (189 dpi) only three of ten animals had cleared the infection.

The *uxt1*Δ *uxt2*Δ mutant induced a generally subdued pulmonary immune response compared to that induced by WT cryptococci. Notably, levels of IL-12p40, a cytokine normally required for inducing a protective response against *C. neoformans* (33), were elevated at 63 dpi and beyond (Fig. 1F); this may facilitate subsequent resolution of the *uxt1*Δ *uxt2*Δ infection. The rise in IL-12p40 could reflect an increase in either IL-23 or IL12 (34), which stimulate Th17 and Th1 type responses, respectively. Since we observed a trend towards increased IL-17 at 63 dpi, but no difference in IFN-γ levels, IL-12p40 was more likely associated with a Th17 response.

In terms of overall lung histology, the early host response of mutant-infected mice was limited. By 128 dpi it had become more organized, with all fungi apparently contained within inflammatory structures, although there was a slight degradation of fungal control by the last time point imaged (189 dpi). We specifically quantitated the temporal progression for iBALT and found that B-cell follicles at the earliest time point of mutant infections (15 dpi) were sparse and small compared to 15 dpi wild-type iBALT, with larger T cell cuffs and smaller areas of CXCL13 staining. By 126 dpi, however, their follicle size and CXCL3 staining had surpassed WT levels, although these parameters regressed somewhat by 189 dpi. The development of iBALT, therefore, paralleled the ability of the host immune response to physically contain cryptococci within inflammatory structures in the lung.

DCs reacted more strongly to *uxt1*Δ *uxt2*Δ than to WT *in vitro*, releasing greater amounts of pro-inflammatory cytokines (Fig. 3B-E), even though the global cytokine response to the mutant was less perturbed (Fig 1D-I). These cytokines may directly or indirectly lead to the increased recruitment of T and B cells in *uxt1*Δ *uxt2*Δ infection (Fig. 2A-C) and perhaps accelerate compartmentalization into compact B cell follicles and discrete T cell areas in iBALT (Fig 5). Such differences in cellular composition and organization, coupled with differences in the local cytokine production, may confer distinct properties on the iBALT induced by WT and *uxt1*Δ *uxt2*Δ, influencing their ability to protect against *C. neoformans*.

We expected that the highly immunostimulatory behavior of *uxt1*Δ *uxt2*Δ *in vitro* reflected the lack of Xyl modifications on its surface glycoconjugates, a consequence of the absence of UDP-Xyl transport into the secretory compartment. We were surprised, therefore, that mutants unable to synthesize UDP-Xyl (*uxs1*Δ), which similarly lack Xyl modifications on secreted glycoconjugates, did not phenocopy this broad and robust DC response. This suggests that Uxs1 itself is required for the strong response seen in the absence of xylosylation in the secretory pathway. It may be that this protein plays additional roles unrelated to glycosylation (35), or that the stimulatory component is dependent on cytosolic UDP-Xyl, rather than UDP-Xyl that has been transported into the secre281 tory pathway.

Unlike encapsulated strains, *C. neoformans* mutants lacking capsule induce the upregulation of multiple genes involved in cytokine responses and enhance antigen processing and presentation by DCs (36). Interestingly, the cytokine levels induced in DC by the *uxt1*Δ *uxt2*Δ mutant were similar to those induced by the acapsular mutant *cap59*Δ (Fig 4A-C). Xyl may thus be critical for the immune suppressive effects normally exerted by capsule material. The mechanisms behind the robust DC activation seen in these two mutants, including the cellular receptors and fungal components responsible, remain to be determined. It is possible that the *uxt1*Δ *uxt2*Δ mutant lacks immune suppressive mechanisms that are present in WT *C. neoformans* infection, and therefore induces robust cytokine responses and consequent recruitment of T and B cells that stimulate iBALT formation and limit early infection.

In this study, we report for the first time that *C. neoformans* infection triggers iBALT formation. To our knowledge, the only other fungus reported to induce iBALT is *Pneumocystis* (37). Notably, the iBALT detected in our experimental infections resembles subpleural nodules that have been observed in cryptococcosis patients (38), although further examination will be necessary to establish the similarities between histological features of inflammatory cell infiltration in mice and humans. Although iBALT formation occurred in mice infected with both mutant and WT fungi, it was inadequate to defend against the overwhelming fungal burden that rapidly develops in a standard murine model of cryptococcal infection. It may be that lower inocula would shift the balance in favor of the host. In any event, the kinetics of iBALT formation and its organization are dependent on virulence factors of the pathogen. Future studies will be needed to fully delineate the molecular host-pathogen interactions that drive iBALT formation following *C. neoformans* infection and the magnitude of the role that iBALT plays; in this regard specific chemokine deficient mice, such as CXCL-13 deficient animals that lack the abil309 ity to induce iBALT structures (39), may be useful.

## Materials And Methods

### Fungal strains

*C. neoformans* strains (Table S1) were grown at 30 °C in YPD medium (1% [wt/vol] yeast extract, 2% [wt/vol] peptone, 2% [wt/vol] dextrose) with shaking (230 rpm) or on YPD agar plates (YPD medium with 2% [wt/vol] agar) supplemented with the following antibiotics as appropriate: 100 μg/ml nourseothricin (NAT; Werner BioAgents) or Geneticin (G418; Invitrogen).

### Mice

C57BL/6J (RRID: IMSR_JAX:000664) and A/JCr (RRID: IMSR_CRL:563) mice were obtained from Jackson Laboratory. *Rag1*-/- (RRID: IMSR_JAX:002216), *TCRβ*-/- (RRID: IMSR_JAX:002118), and μMT mice (RRID: IMSR_JAX:002288) (all on C57BL/6 background) were generously provided by Dr. Michael Diamond and Dr. Wayne Yokoyama (Washington University School of Medicine); breeders were originally purchased from Jackson Laboratory. All mice were 6 to 8 weeks old at the time of infection. A/JCr mice were female; male and female mice were used for the other strains with gender and age matched C57BL/6 controls.

### Ethics statement

All animal protocols were reviewed and approved by the Animal Studies Committee of the Washington University School of Medicine and conducted according to National Insti332 tutes of Health guidelines for housing and care of laboratory animals.

### C. neoformans infection

Fungal strains were cultured overnight (O/N) and diluted to 10^6^ cells/mL in sterile PBS. Mice were intranasally inoculated with a 50 μL aliquot, and then weighed and monitored daily. Infected mice were sacrificed if they lost >20% relative to peak weight or at specified time points. At the time of sacrifice, mice were perfused intracardially with 10 mL sterile PBS, and organs were processed as described below for fungal burden, histology, flow cytometric analysis, and cytokine measurements.

### Organ burden and cytokines

Brain and spleen homogenates were harvested and plated for CFU at the specified time points. 50 μL aliquots of the left lung homogenates (VT = 1 mL) were similarly plated for CFUs. The remaining sample was assayed for pulmonary cytokine levels using the BioPlex Protein Array System (Bio-Rad Laboratories). Briefly, the lung homogenates were mixed with an equal volume of PBS/0.1% Triton 100x/2x protease inhibitor (Pierce EDTAfree protease inhibitor; Thermo Scientific), vortexed for 3 seconds, and clarified by centrifugation (2500 x g, 10 min). Supernatant fractions were then assayed using the BioPlex Pro Mouse Cytokine 23-Plex (Bio-Rad Laboratories) for the presence of IL-1α, IL1β, IL-2, IL-3, IL-4, IL-5, IL-6, IL-9, IL-10, IL-12 (p40), IL-12 (p70), IL-13, IL-17A, granulocyte colony stimulating factor (G-CSF), granulocyte monocyte colony stimulating factor (GM-CSF), interferon-γ (IFN-γ), CXCL1/keratinocyte-derived chemokine (KC), CCL2/monocyte chemotactic protein-1 (MCP-1), CCL3/macrophage inflammatory protein-1α (MIP-1α), CCL4/MIP-1β, CCL5/regulated upon activation normal T cell expressed and secreted (RANTES), and tumor necrosis factor-α (TNF-α).

### Tissue staining and histologic analysis

Mice were perfused intracardially with sterile PBS, and the lungs inflated with 10% formalin. Lung tissue was then fixed O/N in 10% formalin and submitted to the Washington University Developmental Biology Histology Core for paraffin-embedding and sectioning. Immunofluorescence staining for B cells (APC-conjugated rat anti-mouse CD45R/B220, clone RA3-6B2, BD Biosciences, RRID: AB_627078), T cells (CD3-ε, clone M-20, Santa Cruz Biotechnology, RRID: AB_631129), Ig G^+^ plasma cells (FITC-donkey anti-mouse Ig G, Jackson ImmunoResearch Laboratories, RRID: AB_2340796), and CXCL13-producing cells (goat α-mouse CXCL13, AF470, R&D Systems, RRID: AB_355378) was performed as in Ref (37). Images were taken with a Zeiss Axioplan2 microscope and recorded with a Hamamatsu camera. Blinded morphometric analysis of lung structures was performed with the outline tool of Zeiss Axiovision software. Additional sections were stained with Movat’s pentachrome stain by the Washington University Pulmonary Mor371 phology Core.

### Flow cytometric analysis

The right lung of individual mice was enzymatically digested (in 5 mL RPMI with 1 mg/mL collagenase type IV) at 37 °C with shaking (230 rpm) for 30 min, and then sequentially passed through sterile 70 and 40 µm pore nylon strainers (BD Biosciences, San Jose, CA). Red blood cells in the samples were lysed by treatment for 3 min on ice with 5 mL ammonium-chloride-potassium lysing buffer (8.024 g/L NH_4_Cl, 1.001 g/L KHCO_3_, 2.722 mg/L EDTA•Na_2_ 2H_2_O), followed by the addition of 2 volumes of PBS. The remaining cells were pelleted (1000 x g, 5 min, 4 °C), washed twice with PBS, diluted to 10^6^ cells/mL in PBS, and stained with LIVE/DEAD fixable blue dead cell stain (1:1000; Thermo Scientific). Following incubation in the dark for 30 min at 4 °C, cells were washed with PBS and FACS buffer (2% fetal bovine serum in PBS) before resuspension in FACS buffer. Samples were then blocked with CD16/CD32 (1:500; Fc Block™; BD Biosciences, RRID: AB_394656) for 5 min and incubated for 30 min with optimal concentrations of fluorochrome-conjugated antibodies (Table S2) diluted in Brilliant Stain Buffer (BD Biosciences). After three washes with FACS buffer, the cells were fixed in 2% formaldehyde/FACS buffer. For data acquisition, >50,000 events were collected on a BD LSRFortessa X-20 flow cytometer (BD Biosciences), and the data were analyzed with FlowJo V10 (Fig. S1; TreeStar). The absolute number of cells in each leukocyte subset was determined by multiplying the absolute number of CD45^+^ cells by the percentage of cells stained by fluorochrome-labeled antibodies for each cell population analyzed.

### Isolation of bone marrow derived cells

Bone marrow was flushed from the femurs and tibiae of C57BL/6 mice using RPMI. Cells were collected (1000 x g, 5 min, 4°C), resuspended in RPMI, and counted using a hemocytometer. To prepare bone marrow derived dendritic cells (BMDCs), 2 x 10^6^ bone marrow cells were plated in 10 mL R10 medium (10% FBS, 0.4% Penicillin-Streptomycin, 2 mM L-glutamate, 50 μM 2-β-mercaptoethanol in RPMI) supplemented with 1 ng/mL GMCSF, and incubated at 37 °C and 5% CO2. Medium was changed 3 and 6 days after plating, and cells were harvested on day 8. BMDCs were enriched by depletion of BMMs using biotinylated α-F4/80 antibody (eBioscience, RRID: AB_466657) and anti-biotin conjugated magnetic beads (Miltenyi Biotec). The BMDCs in the flow through were positively selected using α-CD11c magnetic beads according to the manufacturer’s protocol (Mil405 tenyi Biotec).

### Dendritic cell assays

To assay the ability of fungal strains to activate BMDCs, *C. neoformans* strains of interest were grown O/N, washed in PBS, and incubated at 65 °C for 15 min to heat kill (HK) the fungi. BMDCs and HK fungi (10^6^ cells of each) were then co-incubated for 24 h, sedimented, and the supernatant fractions transferred to 1.5 mL centrifuge tubes containing 10 μL of 100x protease inhibitor (ThermoScientific). IL-1β, IL-6, and TNF-α levels in supernatants were determined by ELISA according to the manufacturer’s protocol (R&D systems).

### Statistical analysis

Each experiment was performed a minimum of two times. Statistical analyses were conducted using GraphPad Prism version 6.0f (GraphPad Software). All studies comparing two groups were analyzed with a Student’s t-test. Those with three or more groups were compared using an ordinary one-way ANOVA with Tukey’s *post-hoc* test. *p* < 0.05 was considered statistically significant.

## Acknowledgements

We thank members of the Doering laboratory for helpful discussions and assistance with experiments. We also thank Dr. Thomas R. Kozel (University of Nevada School of Medicine) for anti-GXM monoclonal antibody, and Dr. Michael Diamond and Dr. Wayne Yoko427 yama for immune deficient mouse lines.

This work was funded by National Institutes of Health grants R21 AI109623 and R01 AI102882 to TLD. LXL was partly supported by a National Research Science Award (T32 GM007200), a Sondra Schlesinger Graduate Fellowship (Washington University St. Louis Microbiology Department), and a National Institute of Allergy and Infectious Diseases fellowship (F30 AI120339). CRH was partly funded by a National Institute of Allergy and Infectious Diseases training grant (T32 AI007172). SAK and JRM were supported by NIH grant R01 AI111914. The funders had no role in study design, data collection and inter436 pretation, or the decision to submit the work for publication.

## Author Contributions

Conceptualization, LXL, CRH, SAK, and TLD; Methodology, LXL, CRH, JRM, SAK, TLD; Investigation, LXL, CRH, and JRM; Writing – Original Draft, LXL and CRH; Writing – Review & Editing, CRH, JRM, SAK, and TLD; Funding Acquisition, LXL, SAK and TLD; Re442 sources, SAK and TLD; Supervision, SAK and TLD.

## Supplementary Material

**Supplemental Figure 1.**
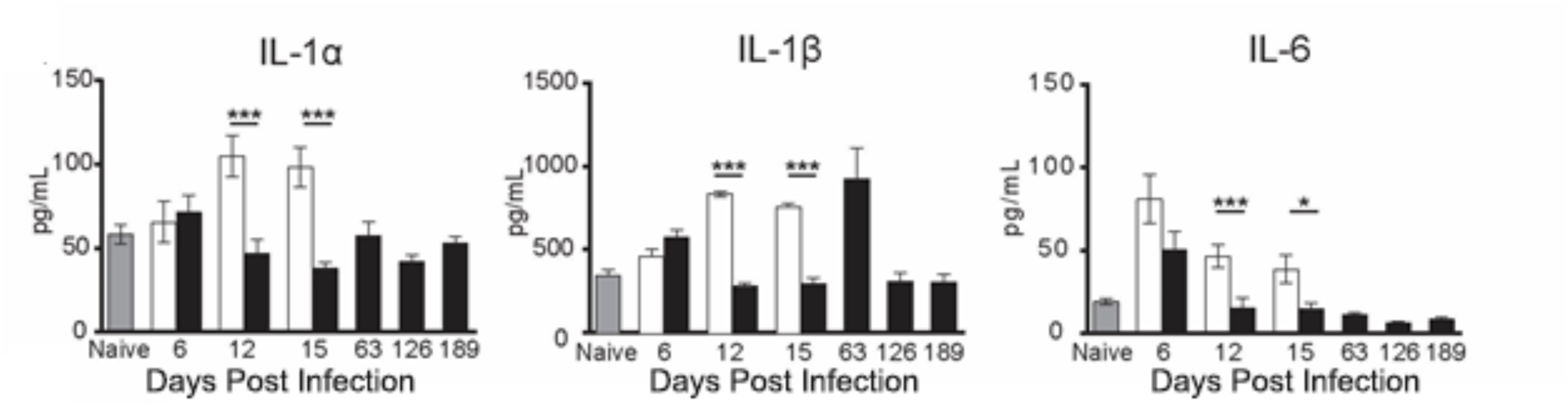
Pro-inflammatory cytokines in lung homogenates at the indicated dpi (grey, naïve; white, WT; black, *uxt1*Δ *uxt2*Δ). Results shown are for individual mice from two independent experiments (n = 5 per group per experiment) and are plotted with the mean ± SEM. *, *p* < 0.05; **, *p* < 0.01; ***, *p <* 0.005 by Student’s t*-*test.

**Supplemental Figure 2.**
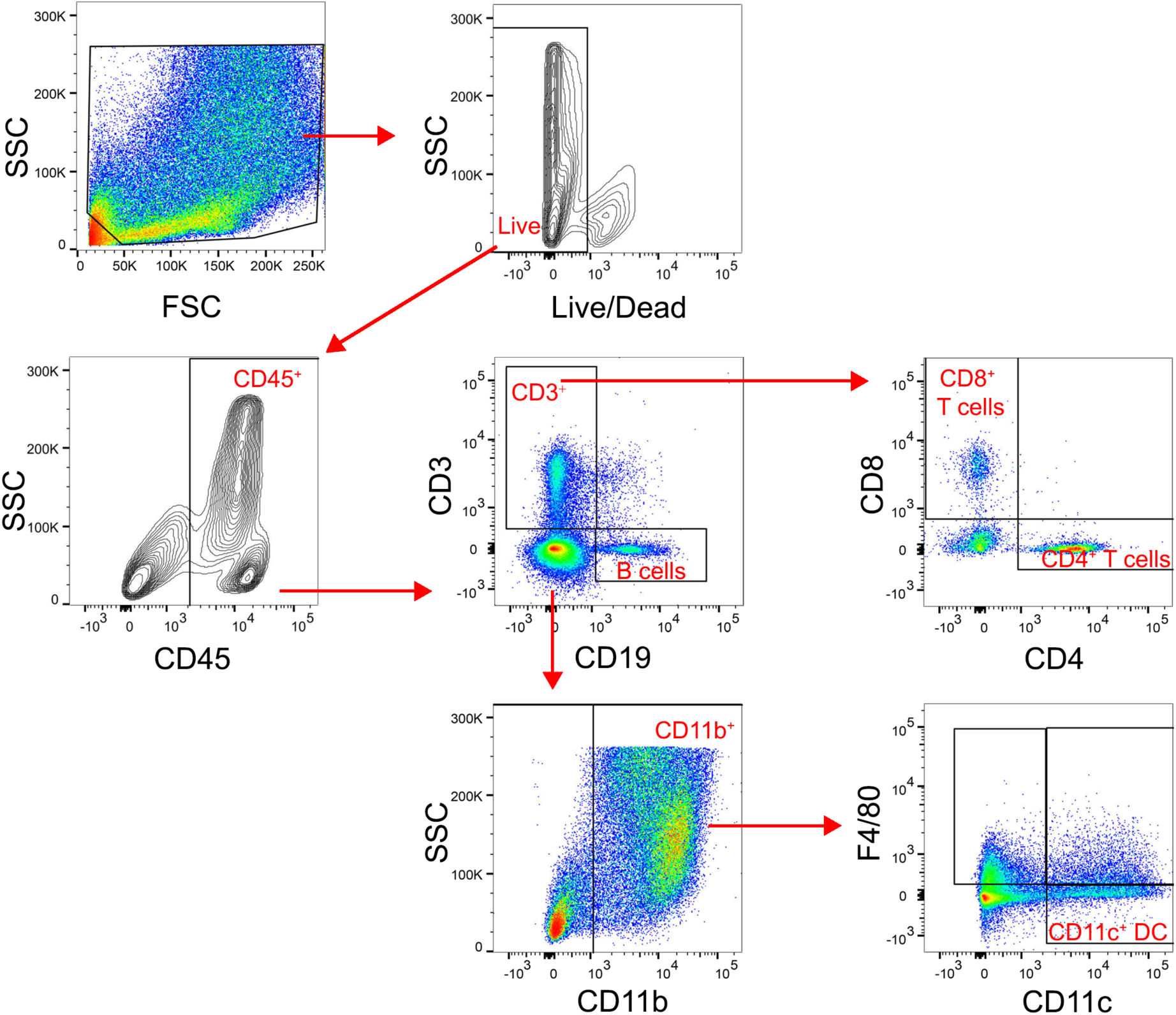
Example of the multi-color flow cytometry gating strategy used to enumerate lung infiltrating immune cells.

**Table S1.** *C. neoformans* strains utilized in these studies.

| *C. neoformans* strain<sup>a</sup> | Origin |
| --- | --- |
| KN99α | (Nielsen et al., 2003) |
| *uxt1*Δ | (Li et al., 2018) |
| *UXT1* | (Li et al., 2018) |
| *uxt2*Δ | (Li et al., 2018) |
| *UXT2* | (Li et al., 2018) |
| *uxt1*Δ *uxt2*Δ | (Li et al., 2018) |
| *uxs1*Δ | (Gish et al., 2016) |
| *cap59*Δ | (Moyrand and Janbon, 2004) |

<sup>a</sup> All mutant strains are derived from KN99α

**Table S2.** Antibodies for flow analysis.

| Antigen | Clone | Fluorophore | Dilution | Company |
| --- | --- | --- | --- | --- |
| CD3 | 17-A2 | APC-Cy7 | 1:250 | BioLegend |
| CD4 | GK1.5 | BUV737 | 1:500 | BD Biosciences |
| CD8a | 53-6.7 | BV650 | 1:125 | BD Biosciences |
| CD11b | M1/70 | BV510 | 1:250 | BD Biosciences |
| CD11c | N418 | PE | 1:125 | BD Biosciences |
| CD1/CD32 Fc Block | 2.4G2 | (not applicable) | 1:500 | BD Biosciences |
| CD19 | 1D3 | BV786 | 1:125 | BD Biosciences |
| CD45 | 30-F11 | Pacific Blue | 1:250 | BioLegend |
| F4/80 | BM8 | APC | 1:250 | Thermo |

